# optSelect: using agent-based modeling and binary PSO techniques for ensemble feature selection and stability assessment

**DOI:** 10.1101/206185

**Authors:** Karan Uppal, Eva K. Lee

**Affiliations:** Center for Operations Research in Medicine and HealthCare, Georgia Institute of Technology, Atlanta, GA, 30332.; NSF I/UCRC Center for Health Organization Transformation, Georgia Institute of Technology, Atlanta, GA, 30332.; School of Industrial and Systems Engineering, Georgia Institute of Technology, Atlanta, GA, 30332.; School of Biology, Georgia Institute of Technology, Atlanta, GA, 30332.

## Abstract

**Motivation:** Recent studies have shown that the ensemble feature selection approaches are essential for generating robust classifiers. Existing methods for aggregating feature lists from different methods require use of arbitrary thresholds for selecting the top ranked features and do not account for classification accuracy while selecting the optimal set. Here we present a two-stage ensemble feature selection framework for finding the optimal set of features without compromising on classification accuracy.

**Methods and Results:** We present herein optSelect, a multi agent-based stochastic optimization approach for nested ensemble feature selection. Stage one involves function perturbation, where ranked list of features are generated using different methods and stage two involves data perturbation, where feature selection is performed within randomly selected subsets of the training data and the optimal set of features is selected within each set using the optSelect. The agents are assigned to different behavior states and move according to a binary PSO algorithm. A multi-objective fitness function is used to evaluate the classification accuracy of the agents. We evaluate the system performance using the random probe method and using five publicly available microarray datasets. The performance of optSelect is compared with single feature selection techniques and existing aggregation methods. The results show that the optSelect algorithm improves the classification accuracy compared to both individual and existing rank aggregation methods. The algorithm is incorporated into an R package, optSelect.

## 1 INTRODUCTION

Biomarker discovery is a key component of translational biomedical research (Guyon 2003, Lee 2007, Christin 2010). Most omics technologies measure thousands of variables (genes, metabolites, etc.) and often fall under the category of n≪p problems that are prone to model over-fitting due to large number of variables (Guyon 2003, Reunanen 2003, Cawley 2010). The large amount of feature space requires application of variable selection techniques to identify most salient variables and generate robust classifiers. This is crucial for targeted validation experiments, designing follow-up studies, and for diagnostic purposes in clinical practice.

Numerous feature selection algorithms have been developed over the last few decades (Saeys 2007, Ma 2008, Christin 2010). The feature selection methods can be classified as: filter, wrapper, and embedded (Saeys 2007). The filter methods use statistical criteria independent of the classifier to select relevant features and the selected features, e.g. p<0.05, are then used to build/train the model. Methods such as t-test, ANOVA, F-test, Chi-sq test, mutual information, etc. can be classified as filter methods. The wrapper methods use a search strategy to evaluate different combinations of subsets of features and select the best model based on the evaluation using a classifier such as Support Vector Machine (SVM, Vapnik 1998). Different search algorithms such as best subset, genetic algorithms, PSO, etc. can be used for finding the optimal set of features; however these methods are prone to over-fitting (Saeys 2007, Christin 2010). The embedded methods include methods such as recursive feature elimination based on SVM, random forests (RF), Lasso, Elasticnet where the variable selection is built-in (Guyon 2002, Breiman 2001, Tibshirani 1996, Zou 2005). For instance, Lasso is a coefficient shrinkage method and uses a L1 penalty function to assign a value greater than 0 if a feature is relevant, and 0 otherwise (Saeys 2007).

Recent articles have highlighted the importance of aggregating ranking results from multiple methods to achieve a set of stable features that are likely to be reproducible in future studies (Saeys 2008, Boulesteix 2009, Abeel 2010, He 2010). The translation of basic science findings to new interventions has been limited due to irreproducible results (Boulesteix 2009, He 2010, Halsey 2015). The two main reasons of “instable” results: a) data perturbation: inconsistency in selected feature subsets due to sampling variations; b) function perturbation: different rankings of relevance from different methods (Boulesteix 2009, He 2010). Many feature selection approaches use arbitrary rank or significance thresholds to select the number of features without thorough evaluation, which could lead to suboptimal results. Moreover, different algorithms vary differently in performance depending on the distribution of the data and within-class variability (Saeys 2008, Boulesteix 2009). Here we introduce a novel nested ensemble feature selection framework, optSelect, that performs multi-objective optimization using agent-based modeling and binary particle swarm optimization (PSO) techniques to allow aggregation of results from different methods and performs nested feature selection using random subsets of training data to evaluate feature stability (Kennedy and Eberhart 1995, Chuang 2010).

## 2 METHODS

### 2.1 optSelect: An optimization based approach for nested ensemble feature selection and aggregation

A two-stage procedure is used for finding the optimal set of features. In stage one, a ranked list of features is generated using one or more feature selection algorithms selected by the user. The user can select t.test, f.test, recursive feature elimination, random forest, wilcox.test, lasso, elasticnet (Saeys 2007, Christin 2013). The number of features selected impacts the performance of the classifiers and the predictive accuracy (Reunanbam 2003). It is essential to find the optimal set of features. Users have the option to perform sequential backward or forward selection or choose an arbitrary cutoff to select the optimal set of features. Using different ranking criteria allows function perturbation. The different variable selection methods implemented in the CMA package in R are used at this stage (Slawiski 2010). A union of the ranked lists from different methods is used as input for stage two. Users have the option to skip stage one and use all features during the optimization procedure; however some selection criteria is recommended prior to stage two to reduce the computational time and perform ensemble selection (Saeys 2007).

In stage two, the newly developed multi-objective stochastic optimization procedure based on agent-based modeling and binary framework is used to perform data perturbation and aggregation using the results from stage one.

Agent-based models involve three key elements (Macal 2010):

a. Agents and their attributes/behaviors: each agent is assigned to a behavior or rule category, e.g. follows neighbors, moves randomly, etc.
b. Relationships and interactions between agents: the interactions define how the agents influence and cooperate with each other,
c. Interaction with the environment: the environment provides feedback to the agent about their movements (e.g. 10-fold cross-validation accuracy)

A binary PSO algorithm is used to simulate the movement of agents according to their behavior. PSO is a stochastic optimization technique based on the movement and intelligence of swarms developed by James Kennedy and Russell Eberhart in 1995. It comprises of a number of agents/particles that constitute a swarm moving around in the search space looking for the best solution determine based on a fitness evaluation function. The movement of each particle, p_i_, is determined based on a velocity vector, v_i_ and a position vector, x_i_. In binary PSO, the position vector, x_i_, has d dimensions, where d corresponds to the number of variables (genes, chemicals, etc.). And, x_id_={0,l}. This allows application of the binary PSO for feature selection, where a feature is selected if x_id_=1. The velocity and position vectors are updated according to equations (1) and (2).

The velocity of each particle, p_i_, is updated at iteration t+1 according to the equation,

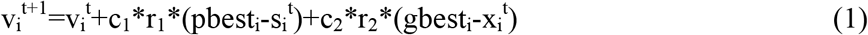

where,

i is the current particle
c_1_ and c_2_ are constant learning factors to control the social influence and global influence, r_1_ and r_2_ are random numbers between [0,1] interval,
pbest is the best position of the particle has experienced based on the fitness function,
gbest is the best position experienced by any particle in the swarm based on the fitness function, and
x_i_ is the position in iteration t.

The velocity of the particle is restricted to be in the interval [-6,6], The position is updated according to (2) and (3) in binary PSO based on a sigmoid transformation, S, of the velocity (Figure 1), for d in 1 to number of variables:

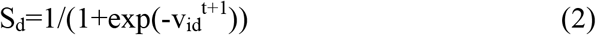

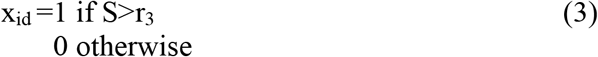

where

x_id_^t^ is the position of the i^th^ agent at time t in dimension d (gene, chemical, etc.),
v_id_^t+1^ is the updated velocity of the i^th^ agent at iteration t+1 in dimension d,
S_id_ is the sigmoid function with values between [0,1] interval for dimension d,
r_3_ is a random number between [0,1] interval

In the modified binary PSO proposed in this work, users can provide a weight vector to bias the selection process based on expert knowledge or from literature. Studies have shown that incorporating prior knowledge can improve the classification accuracy (He 2010). The random number in (3) is replaced by the weight of the feature [0 to 1] to bias the selection process.

**Figure 1.**
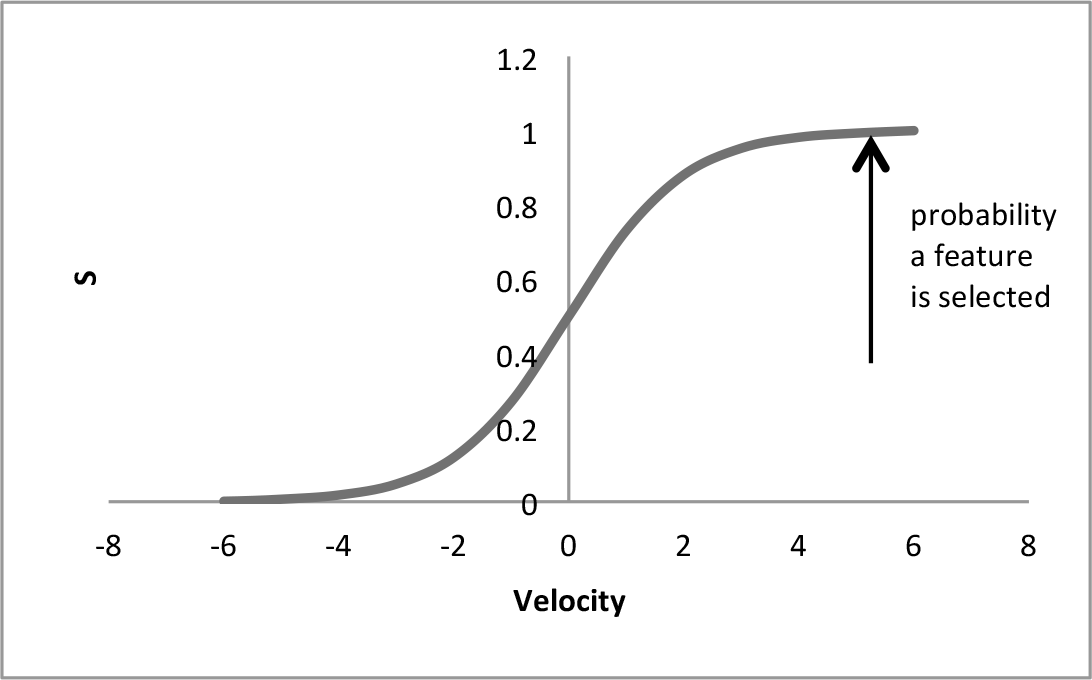
Relationship between velocity, sigmoid function, and probability of a feature being selected. The likelihood of a feature being selected increases as the velocity approaches −6 and S approaches 1.

In most existing versions of the binary PSO algorithm, all particles behave uniformly and generally follow the fully connected topology where each particle is connected to every other particle (Chuang 2008). Each agent is assigned to one of the four behavioral states {C=Confusion, S=Self-influenced, N=Influenced by nearest neighbors, G=Influenced by swarm} based on the crowd model (Wu 2014). The behavior of the current particle determines which nodes in the swarm network are chosen for interaction and updating velocities as described below:

a. Behavior=N, follows neighbors

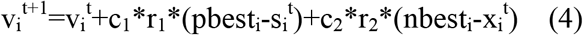

where nbest is the centroid (75^th^ percentile) of the k nearest neighbors (default k=3),
b. Behavior=G: follows global leader

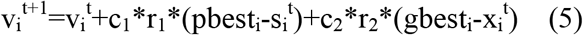

where gbest is the global best position in the swarm,
c. Behavior=C: moves randomly

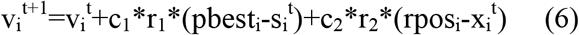

where rpos is a random position vector,
d. Behavior=S: only self-influenced

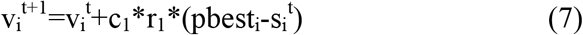

The behaviors are updated on regular intervals to prevent the population from getting stuck in local optima (user-defined parameter; default: 5 iterations). The fitness of each particle is evaluated using a nested cross-validation procedure and support vector machine (SVM) classifier as shown in Figure 2 (Vapnik 1998). The algorithm uses the internal cross-validation scheme that performs variable selection based on the training set, trains_k_, in the m-fold scheme and uses the left-out subset, valid_m_, for evaluating the performance of the model. The internal cross-validation scheme is shown to out-perform the commonly used external cross-validation scheme where the model evaluation is performed after the selection using all samples (Cawley 2010). The search process is terminated when the distance between the centroid of the entire population and the global best agent is less than or equal to 2.

### 2.2 Evaluation experiments

Two experiments were performed to evaluate the performance of the newly developed framework. In the first experiment, a random probe evaluation was performed using the Iris dataset (Fisher 1950). The original dataset has 4 real features and 150 instances belonging to 3 classes of the iris plant. Each real feature was scale normalized (*µ* = 0, *σ* = 1) and 36 random features with similar distribution were included in the dataset. The instances were divided into 60% train (90; 30 per class) and 40% test (60; 20 per class). The aim was to evaluate the ability of the stage two of the optSelect algorithm to identify real features. In the second experiment, five publicly available microarray datasets: Leukemia (7129 genes, train samples=38, test samples=34, 2 classes; Golub 1999), SRBCT (2308 genes, train samples=63, test samples=25, 4 classes; Khan 2001), Prostate cancer (6033 genes, train samples=61, test samples=41, 2 classes; Singh 2002), MAQCII-ER and MAQCII-PCR (22284 genes, train samples=130, test samples=100, 2 classes) (Popovici 2010). Limma, rfe-SVM, Lasso, Elasticnet, F-test methods implemented in the CMA R package were used for feature selection in stage one. The two rank aggregation methods implemented in the rankAggreg R package, rankAggreg-Monte Carlo and rankAggreg-GA, that optimize the list of ranked features based on the Spearman’s footrule were used for comparison in stage two. The evaluation process was repeated using top 5 and top 15 features from stage one. The final performance of different methods is evaluated using an independent blinded test set that was not seen by the optSelect algorithm during the model building stage. The balanced accuracy, average class-wise prediction accuracy, was used for evaluation purpose (Dreyfus 2006).

**Table 1.** Using top 5 ranked features in Stage 1. The balanced accuracies for the independent test sets are reported here.

|  | Method\Dataset | Leukemia | SRBCT | MaqcII_ER | MaqcII_PCR | Prostate |
| --- | --- | --- | --- | --- | --- | --- |
| Stage 1 | Limma | 0.89 | 0.94 | 0.87 | 0.7 | 0.77 |
|  | rfe-SVM | 0.94 | 0.92 | 0.88 | 0.66 | 0.77 |
|  | Lasso | 0.94 | 0.9 | 0.87 | 0.62 | 0.75 |
|  | Elasticnet | 0.89 | 0.94 | 0.87 | 0.72 | 0.77 |
|  | F.test | 0.93 | 0.94 | 0.87 | 0.66 | 0.77 |
| Stage2 |  |  |  |  |  |  |
| (Aggregation) | rankAggreg-monte carlo | 0.89 | 0.92 | 0.87 | 0.74 | 0.77 |
|  | rankAggreg-GA | 0.89 | 0.94 | 0.87 | 0.64 | 0.77 |
|  | **optSelect** | 0.96 | 1 | 0.87 | 0.72 | 0.77 |

## 3 RESULTS

For experiment 1, 3 out of 4 real features selected in 8 out of 10 folds. 2 were selected in all 10 folds. All random features were discarded. Both train 10-fold cross-validation accuracy and balanced accuracy for the test set were 95%.

The results for experiment 2 are summarized in tables 1 and 2. Table 1 shows the test set balanced accuracies for the test sets from the five datasets using top 5 ranked features selected by different methods (limm, rfe-SVM, Lasso, Elasticnet, F.test) in stage 1 followed by different aggregation methods in Stage

**Figure 2.**
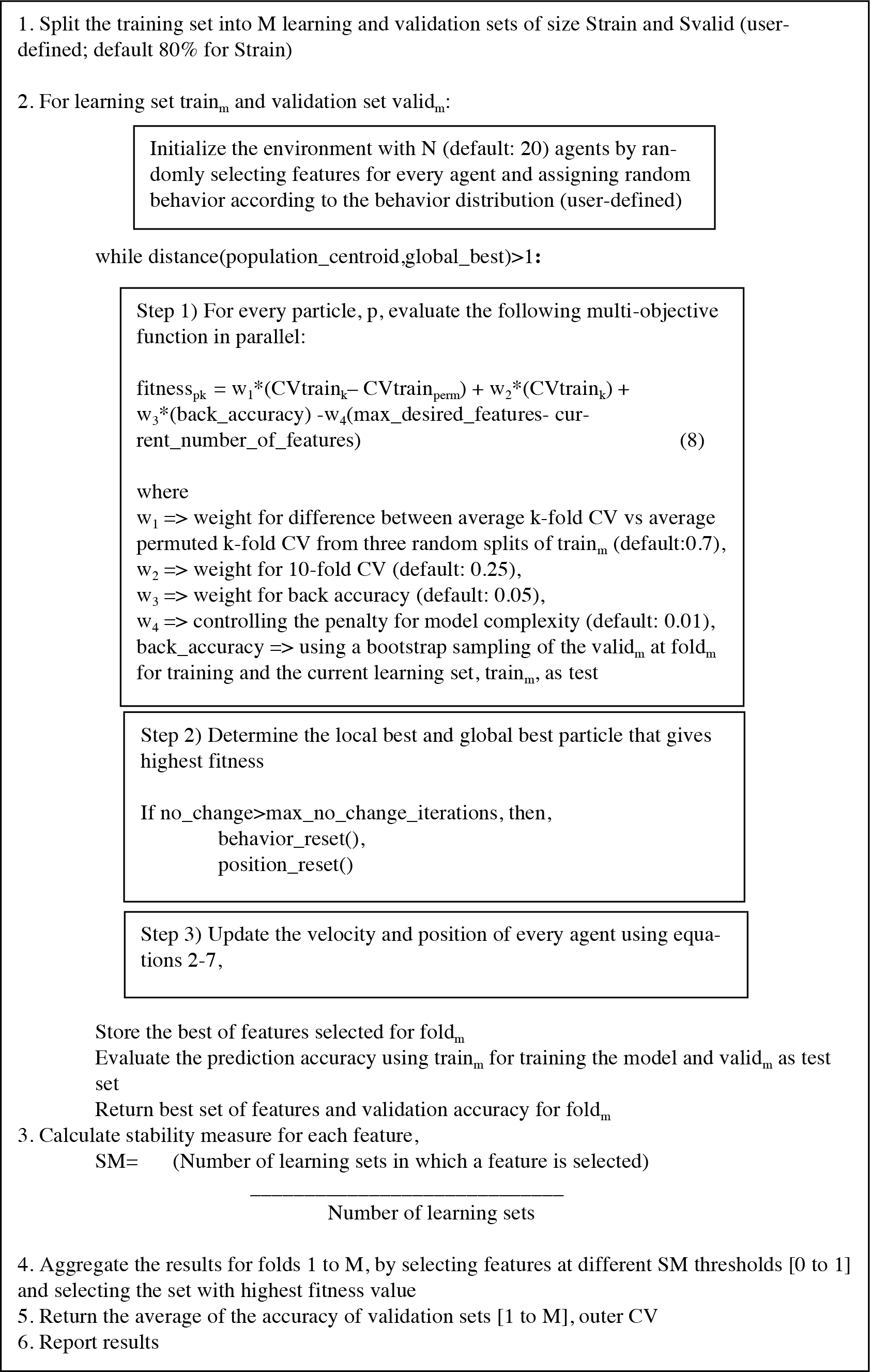
optSelect algorithm for nested feature selection

The results show that the ensemble approach improves the classification results as compared to individual methods. For both Leukemia and SRBCT test sets, the optSelect algorithm achieved highest accuracy compared to both individual methods in stage 1 and existing aggregation techniques. For the other three datasets, the optSelect algorithm gave similar performance as compared to individual and existing methods.

Table 2 summarizes the balanced accuracy results for the five datasets using the top 15 ranked features selected by different methods (limm, rfe-SVM, Lasso, Elasticnet, F.test) in Stage 1 followed by different aggregation methods in Stage 2. As discussed earlier, the arbitrary thresholds for selecting top ranked features does not guarantee reproducibility of results on a test set. For instance, the performance of rfe-SVM and Elasticnet degrades by 28% for the SRBCT dataset by increasing the number of selected features to 15. On the contrary, almost all methods showed improvements in accuracy for the Leukemia dataset by using greater number of features. Overall, the aggregation stage improved the classification accuracy with the optSelect algorithm performing comparably or better in almost all cases.

**Table 2a.**
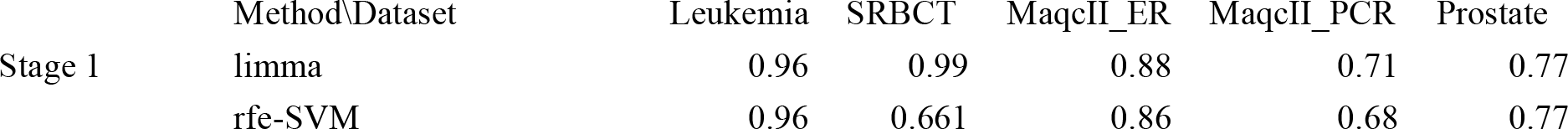
Using top 15 ranked features in Stage 1. The balanced accuracies for the independent test sets are reported here.

**Table 2b.** Using top 15 ranked features in Stage 1. The balanced accuracies for the independent test sets are reported here.

|  |  |  |  |  |  |  |
| --- | --- | --- | --- | --- | --- | --- |
|  | Lasso | 0.96 | 1 | 0.87 | 0.64 | 0.77 |
|  | Elasticnet | 0.96 | 0.66 | 0.88 | 0.68 | 0.77 |
|  | F.test | 0.96 | 0.986 | 0.88 | 0.66 | 0.77 |
| Stage2 |  |  |  |  |  |  |
| (Aggregation) | rankAggreg-monte carlo | 0.94 | 1 | 0.88 | 0.75 | 0.77 |
|  | rankAggreg-GA | 0.96 | 0.96 | 0.87 | 0.73 | 0.77 |
|  | **optSelect** | 0.96 | 1 | 0.9 | 0.73 | 0.83 |

## 4 DISCUSSION

In recent years, various articles have raised the issue of feature instability and irreproducibility that has hindered the translation of results from basic science to clinical domain (Boulesteix 2009, Abeel 2010, He 2010, Halsley 2015). Several methods for rank aggregation have been proposed (Saeys 2008). However, most of these methods use criteria independent of the classification accuracy for aggregating the results and require selection of an arbitrary threshold for top k features to determine the overlap. This could result in degradation of classification performance on an independent or unseen data set.

Here we propose a novel optimization based nested ensemble feature selection framework, optSelect, which addresses this problem by using a two-stage approach for aggregating the results. Stage one involves selection of top ranked features using different methods. The top list of features from different methods is merged and used as input for a multi agent-based optimization procedure to find the most stable set of features with good classification accuracy. The “optimal” set is determined using a multiobjective fitness function as described in Methods. The algorithm incorporates the concepts of function perturbation and data perturbation to select the most optimal and stable set of features (Boluestrix 2009, He 2010). Additionally, the algorithm is designed to prevent agents from getting stuck in local optima. Each agent interacts with other agents based on their behavior state {confusion, follows neighbors, follows global leader, self-influenced}. Search for optimal solution terminates when the global best and the centroid of the population converge, which is determined using the Euclidean distance.

The performance evaluation of the multi agent-based optimization procedure on the random probe experiment (Experiment 1) shows that the newly proposed behavior based search algorithm combined with the multi-objective classification based fitness function allows detection of relevant features even when majority of the features are randomly generate. Evaluation results for the five gene expression datasets (Experiment 2) show that the newly proposed optSelect framework allows aggregation of results from different methods without compromising for the classification accuracy. The results also highlight the dramatic changes in classification accuracies as a result of arbitrary thresholds for selecting top features. Furthermore, the performance of different independent feature selection techniques varies across different datasets. On the contrary, the optSelect algorithm performed consistently across all datasets and improved the classification results on independent test sets in most cases. The output includes outer CV estimates, optimal set of features, and stability measures for each features. Figure 3 shows the stabil-ity measures of the 6 features from the optimal set selected by optSelect for the MAQCII-ER dataset, where the samples were classified as ER+ve vs ER-ve. The most stable feature that was reproducibly selected in all folds in the nested feature selection was estrogen receptor 1.

**Figure 3.**
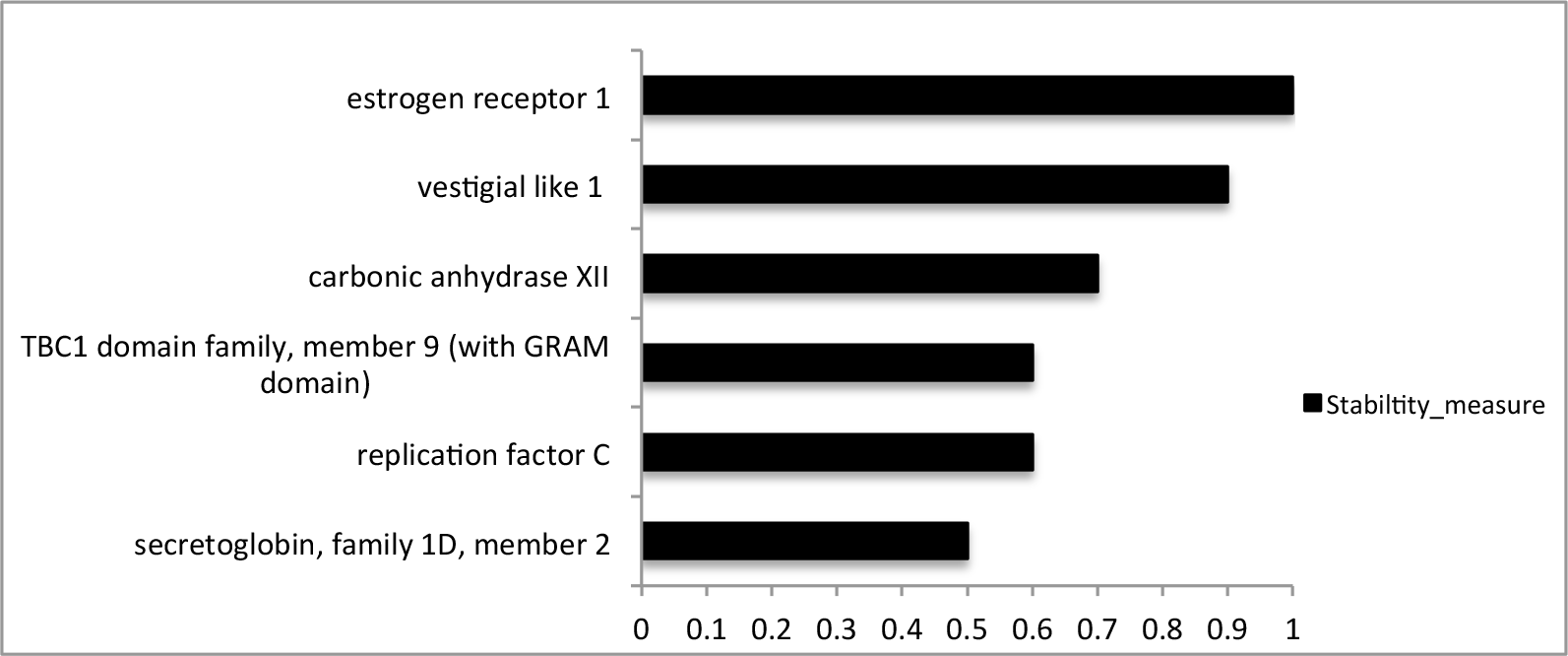
Stability measures for features in the optimal set selected by optSelect.

**Limitations and future work:** The current study did not assess the effect of using data normalization methods and classifiers on classification accuracy. Escalente et al. have recently showed application of PSO for full model selection (pre-processing methods, feature selection, and classification algorithms). Future work will focus on extending the current framework to full model selection.

## 5 CONCLUSION

We have developed a novel multi-stage nested ensemble feature selection algorithm, optSelect, which accounts for function perturbation and data perturbation during the feature selection process. The results show that the ensemble framework improves the classification accuracy over independent feature selection methods and existing rank aggregation methods.

